# Constitutive expression of cardiomyocyte Klf9 precipitates metabolic dysfunction and spontaneous heart failure

**DOI:** 10.1101/2025.01.16.633464

**Authors:** Chandni Thakkar, Saleena Alikunju, Aishwarya Venkatasubramanium, Zhi Yang, Nazish Sayed, Maha Abdellatif, Danish Sayed

**Author notes:** To whom correspondence should be addressed: Danish Sayed, Department of Cell Biology and Molecular Medicine, Rutgers New Jersey Medical School, 185 South Orange Avenue, Medical Science Building, G-653/ MSB G-626, Newark, New Jersey 07103. Cardiovascular Institute, Department of Surgery, Division of Vascular Surgery, Stanford University School of Medicine, Stanford, California 94304 (Nazish Sayed).

## Abstract

Metabolic adaptations and flexibility during development and disease play an essential in cardiomyocyte function and survival. We recently reported Glucocorticoid receptor (GR)-Krüppel-like factor 9 (Klf9) axis in mediating metabolic adaptations in cardiomyocytes stimulated with Dexamethasone. Klf9 expression decreases in hypertrophic and failing hearts, suggesting its importance in cardiac homeostasis and its potential contribution to dysfunction under pressure overload. Genome wide Klf9 occupancy in adult hearts revealed 2,242 genes directly associated with Klf9, with enrichment in metabolic pathways, autophagy, ubiquitin-mediated proteolysis, and cellular senescence. We generated and characterized a conditional cardiac specific Klf9 knock-In (Klf9KI) mice, which developed progressive cardiac hypertrophy, cardiac dysfunction and cardiac failure by 8wks of age, with mortality by 12-14wks. RNA-seq analysis at 1wk, 4wks, and 8wks showed stage-specific transcriptional changes. At 1 week, 64.81% of differentially expressed genes were downregulated, aligning with Klf9’s predicted role as a transcriptional repressor. At 4wks and 8wks, more genes were upregulated, suggesting more of secondary targets in response to cardiac phenotype. KEGG pathway analysis showed dysregulation in lipid, carbohydrate and glutathione metabolism, transcriptional regulation, apoptosis, and innate immunity. Untargeted Metabolomics at 4wks identified significant alterations in tissue metabolite levels, particularly in pathways involving fatty acid metabolism, amino acids, and glucose, correlating with transcriptome data. Mitochondrial function assays revealed progressive dysregulation. At 2 weeks, complex I activity was significantly reduced, while complex II and IV activities were partially preserved. By 4 weeks, all measured respiratory complexes showed significant declines, consistent with decline in mitochondrial function. These mitochondrial deficits preceded overt cardiac dysfunction and likely contributed to the development of hypertrophy and failure. In conclusion, constitutive Klf9 overexpression disrupts transcriptional and metabolic homeostasis, driving progressive hypertrophy, cardiac dysfunction, and failure.

## Introduction

Krüppel-like factors (Klf) have been shown to regulate various cellular processes, including metabolism, differentiation, proliferation with implications in development and diseases of cardiovascular, nervous, respiratory and immune systems (1, 2). We recently reported an essential role of Klf9 in transcriptional control of metabolic genes, and a GR-Klf9 axis in metabolic adaptation in response to Dexamethasone stimulation in cardiomyocytes (3). Klf9, with enriched expression in the heart has been shown to be downregulated in rodents and patients with cardiomyopathy (4), further, transcriptome analysis from human failing and control (non-failing) hearts shows downregulation of Klf9 transcript with both HFpEF and HFrEF (5), suggesting that Klf9 expression status and function correlates with cardiac pathogenesis. Here we report the characterization of cardiac-specific (αMHC) Klf9 knock-in (Klf9KI) mice that develops progressive postnatal cardiac hypertrophy and dysfunction followed by spontaneous heart failure by 8wks, and death by 12-14wks of age.

## Results

Klf9 expression decreases in hearts undergoing pressure overload induced cardiac hypertrophy (Fig S2), these observations have also been recently shown in mouse hearts undergoing Angiotensin II induced cardiac hypertrophy and failure (4). A reduced RNA pol II recruitment and distribution across the Klf9 gene structure in hypertrophy hearts compared to sham (6) indicates a decrease in transcription and expression of Klf9 (Fig S2A), which corresponds with decrease in Klf9 transcript abundance in these hearts undergoing hypertrophy (Fig S1B and S1C). Klf9 expression levels decrease prior to decrease in cardiac dysfunction, starting 7day post TAC. These data suggest that Klf9 plays an essential role in cellular homeostasis in adult hearts and a decrease with pressure overload could be contributing to the progression of cardiac dysfunction and heart failure.

To that end, we generated conditional cardiac-specific Klf9 knock-in (Klf9KI) mice (Fig S1A), with 3-4-fold increase in Klf9 expression when mated with αMHC-Cre expressing mice (Fig S1B and C). Interestingly, these mice developed spontaneous progressive hypertrophy by 4wks of age (Fig 1A, 1B and 1C), followed by cardiac dysfunction and failure by 8wks (Fig 1D and 1E). Klf9KI mice showed early mortality by 12-14wks, in both male and female mice (Fig 1F), suggesting that constitutive increase in cardiac Klf9 expression could be detrimental to the postnatal cardiac development and function. Next, to determine the transcriptional targets of cardiac Klf9, we performed Klf9-ChIP-seq in mouse hearts that identified 2242 genes that showed Klf9 genomic association (Fig 2A and 2C, supplementary Excel sheet), with 71.08% of the genes with intervals within 500bp of the transcription start site (Fig 2B). KEGG pathway analysis of these genes showed maximum genes involved in metabolic pathways, along with top five categories including endocytosis, ubiquitin mediated proteolysis, autophagy-animal, focal adhesion and cellular senescence (Fig 2D and S3). Further characterization of the genes from the metabolic pathways, revealed genes involved in biosynthesis of cofactors, carbon metabolism, metabolite metabolism, fatty acid metabolism, glycolysis/gluconeogenesis, glycerophospholipids, glucagon signaling, starch & sucrose metabolism, lipoic metabolism and Glutathione metabolism (Fig 2E). These data from adult hearts align with our published data from neonatal cardiomyocytes stimulated with dexamethasone (Dex, synthetic glucocorticoid, with high affinity for Glucocorticoid receptor), showing role of GR-Klf9 axis in cardiomyocyte metabolic adaptations (3).

**Figure 1.**
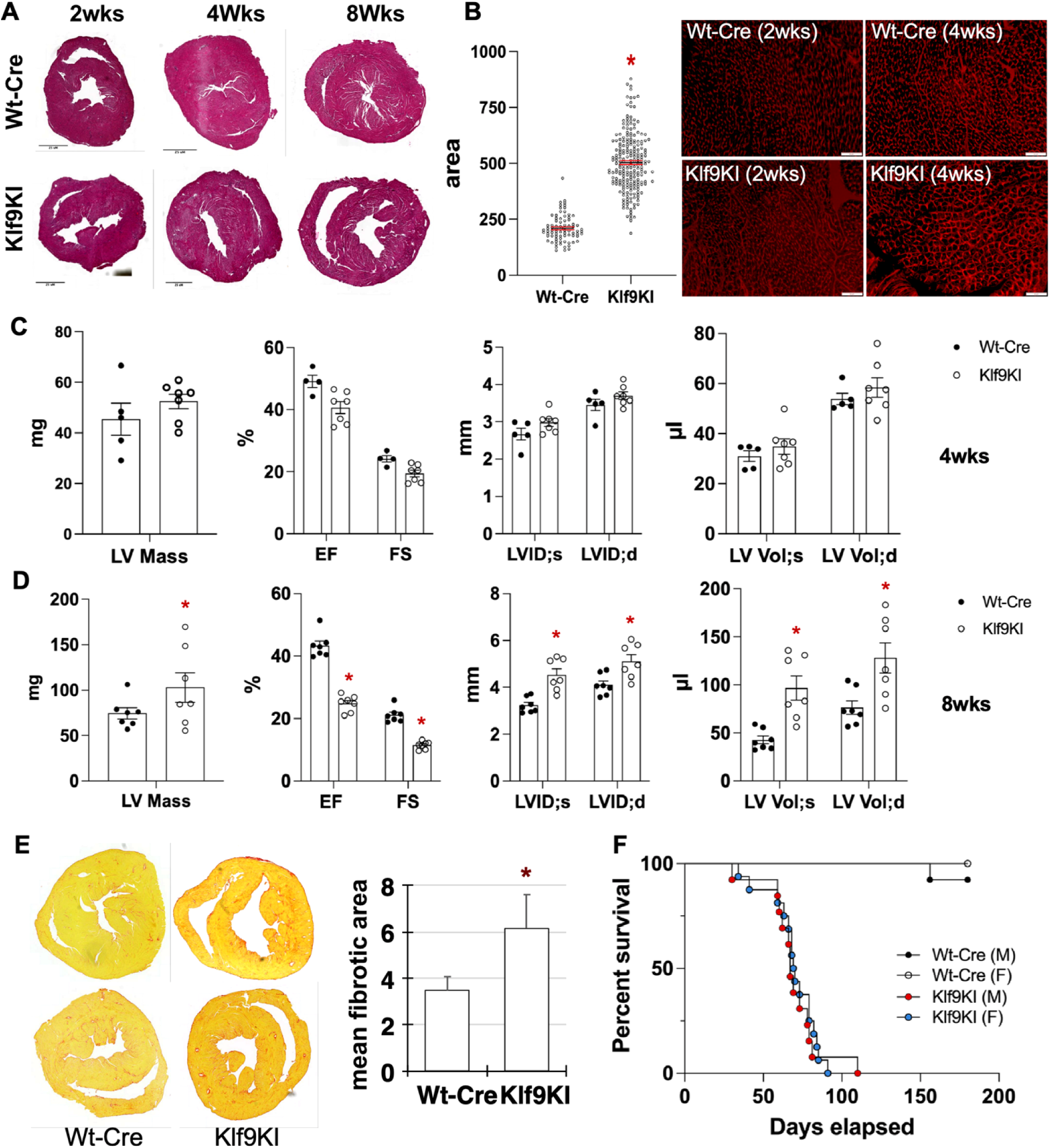
Physical, morphological and functional characterization of Klf9KI and Wt-Cre mice. **A.** H & E staining of cross sections of hearts from 2wks, 4wks, and 8wks old Wt-Cre and Klf9KI mice indicating progressive hypertrophy. **B.** Graph represents the cross-sectional area of in individual cardiomyocytes in sections from 4wks hearts of Wt-Cre and Klf9KI mice. Representative images of the WGA staining from 2wks and 4wks are shown. **C.** and **D.** Graphs represents left ventricular mass (LV mass), percent ejection fraction (%EF), percent fractional shortening (%FS), left ventricular internal dimensions during systole and diastole (LVID;s, LVID;d), left ventricular volume with systole and diastole (LV Vol;s and LV Vol;d) in Wt-Cre and Kltf9KI mice, as measured by 2D echocardiography. **E.** Representative images of PASR staining in Wt-Cre and Klf9KI mice showing the cardiac fibrosis. Graph represents mean fibrotic area as measured using image J. **F**. Survival curve for Wt-Cre and Klf9KI mice (males and females) generated on Prism 10 on cohorts (n=12 females and 14males Klf9KI mice with their respective Wt-Cre controls). For all graphs, error bars represent SEM, * is p<0.05 compared to respective Wt-Cre, n=3-7 independent hearts.

**Figure 2.**
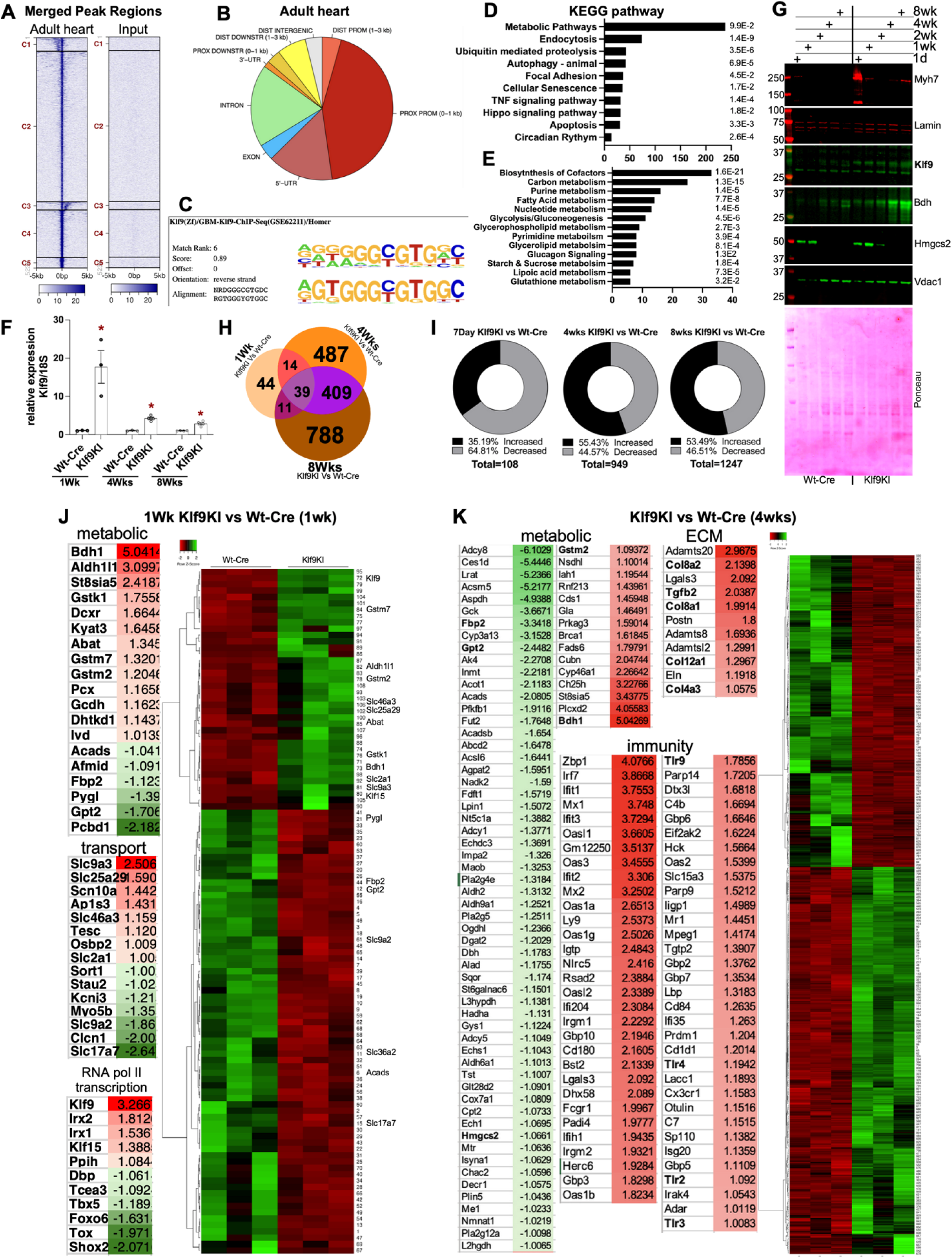
Genomic targets of cardiac Klf9 and differential transcriptome in Klf9KI versus Wt-Cre hearts. **A.** Heatmap showing tag distribution for Klf9 across merged peak regions (values in z-axis/color, active regions in y-axis) in sham hearts and Input control. The data is presented in 5 clusters (default), C1 to C5 and sorted. **B.** Pie chart shows the location of the Klf9 peaks relative to genomic annotations in hearts of adult mice. **C.** Enriched motif identified using Homer Motif analysis using the 200 bp surrounding the summit of the top 2,500 peaks (based on MACS2 p-values). **D** Graphs lists the number of genes in the functional groups as categorized by KEGG pathway analysis, which show Klf9 association in adult hearts. KEGG pathway using DAVID was performed on 2242 genes which show Klf9 genomic binding (Maxtags≥50). **E.** Graph list the number of genes in the sub metabolic groups upon KEGG pathway analysis of the 238 genes from the metabolic pathways from 2D. **F.** Graph represents the relative Klf9 expression at 1wk, 4wks and 8wks old Klf9KI hearts compared to Wt-Cre. Error bars represent SEM, * is p<0.05 compared to respective Wt-Cre, n=3-4. **G.** Western blotting of total protein lysate from 1day, 1wk, 2wks, 4wks and 8wks old Wt-Cre and Klf9KI mouse hearts probed for the indicated proteins. Lamin A/C and Ponceau stating dhow for loading controls. **H.** Venn diagram shows number of genes that are differentially regulated between Wt-Cre and Klf9KI mice at 1wk, 4wks, and 8wks age (|log2FC|≥1), n =3 for each time point with their respective controls. **I.** Donut Pie charts show the number of differentially regulated genes, percent of genes that are upregulated and downregulated in 1wk, 4wks and 8wks old Klf9KI versus Wt-Cre mice. **J.** Table lists the differentially expressed genes from selected pathways in 1wk old KLf9KI and WT-Cre hearts. Heatmap represents levels of the 108 differentially expressed genes in these hearts. **K.** Table lists the differentially expressed genes from selected pathways in 4wks old KLf9KI and WT-Cre hearts. Heatmap represents 949 differentially expressed genes in these hearts.

These data indicates that Klf9 regulates metabolic adaptations in neonatal and adult cardiomyocytes, and under external stimulus could be involved in mediating metabolic flexibility and homeostasis to changing physiological variations, like food intake, circadian cycle, and cardiac stress like pressure overload.

Further, to determine the effects of constitutive expression of Klf9 on the cardiac transcriptome, we performed total RNAseq on 1wk, 4wks and 8wks Klf9KI and Wt-Cre hearts (2F and 2G). As expected, we observed differential regulation in increasing number for genes between 1wk, 4wk and 8wks (Fig 2H). Interestingly, in accordance with the transcription repressive predicted function of Klf9 (1), we observed a decrease in 64.81% of the differentially expressed genes, while dynamics changed in 4wks and 8wks hearts, with more upregulated genes in Klf9KI mice compared to Wt-Cre (Fig 2I and S4A), suggesting secondary differential expression in response to developing hypertrophy and onset of heart failure. Functional annotation and KEGG pathway analysis using DAVID identified these as genes of the metabolic pathway on the top of the list in 7day Klf9KI versus Wt-Cre hearts, with increase in genes involved in Lipid metabolism (for e.g. Bdh1, St8sia5) and glutathione (Gstk1, Gstm7, Gstm2) pathways while a decrease is observed in genes involved in carbohydrate and glycogen (Fbp2, Pygl, Phkg1) metabolism. In addition, genes involved in transport (Slc9a3, Sort1) and RNA pol II dependent transcription (Klf15, Foxo6, Tbx5) were also differentially regulated in Klf9KI mice compared to Wt-Cre (Fig 2J and S4E). At 4wks, along with dysregulation of metabolic pathways, we observe an increase in genes involved in apoptosis (Bcl2, Bid), innate immunity (Tlr genes) and extracellular matrix (Col8a1, Tgfb2) (Fig 2K and S3F), coinciding with the onset of cardiac dysfunction and pathological hypertrophy. To examine the metabolic status in these hearts, we performed untargeted tissue metabolomics in Klf9KI and Wt-Cre hearts at 4weeks of age (Fig 3A), which showed a differential level in several metabolites (Fig 3B) corresponding to dysregulation of metabolites from several metabolic pathways in Klf9KI mice compared to Wt-Cre hearts (Fig 3C). Interestingly, metabolomics data correlated with the differential RNAseq results from 4wks Klf9KI and Wt-Cre hearts, indicating metabolic dysfunction in the hearts, which precedes cardiac dysfunction and onset of failure. By 8wks, when the hearts of Klf9KI mice are showing signs of failure with ventricular dilatation, fibrosis and dysfunction we observed decrease in genes involved in glycogen metabolism, response to Insulin, muscle contraction, transport and lipid metabolism. On the other hand, we observe an increase in proteolysis, apoptosis, extracellular matrix (ECM) and innate immunity genes (Fig S4D). These data show that in adult hearts Klf9 dictates transcriptional expression of genes that are mostly involved in metabolic adaptations and required for cardiomyocyte metabolic flexibility and homeostasis. However, constituent Klf9 levels expression could lead to dysregulated expression of these genes, resulting in metabolic maladaptation with postnatal growth and eventual cardiac dysfunction and progression into failure.

**Figure 3.**
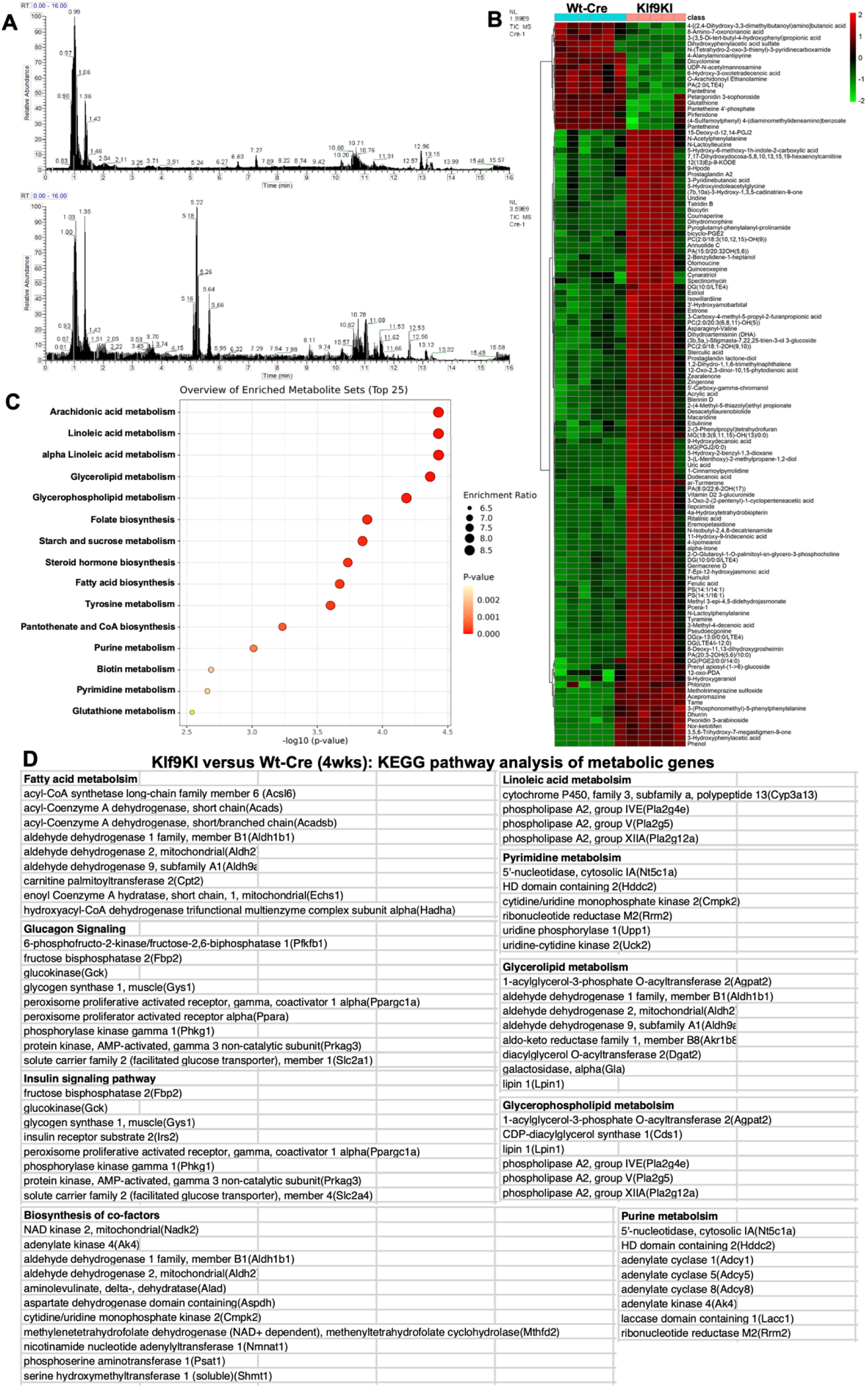
Untargeted tissue metabolomics in Klf9KI and Wt-Cre. **A.** Plot shows the total ions chromatograph (TIC) chromatogram in ESI- (top) and ESI+ (bottom) mode. **B.** Hierarchical cluster analysis of metabolome data from significant metabolites (VIP>1.5, |log2FC|>0.5 and p<0.05) between Wt-Cre and Klf9KI groups. Mean values of metabolite contents from biological replicates of each group are used to calculate the metabolite ratio. Color intensity correlates with degree of increase (red) and decrease (green) relative to the mean metabolite ratio. **C.** Dot plot of network analysis in perturbed metabolites. **D.** Table shows list of genes from selected metabolic pathways from differential RNAseq results in 4wks old Klf9KI and Wt-Cre hearts which correlate with the enriched metabolic pathways from the differential metabolites shown in (C). Metabolomics data include N=6 for the Wt-Cre group and N=5 for the Klf9KI group.

Further, we measured the mitochondrial function and ROS production at 2wks and 4wks in Klf9KI and Wt-Cre mice. A decrease in mitochondrial complex I activity is observed in both age groups of Klf9KI mice, however, complex II and complex IV that correlates with reserve capacity and ATP generation, respectively are partially preserved at 2wks and significantly decreased by 4wks in Klf9KI compared to Wt-Cre mice (Fig 4). Increase in ROS was observed in both 2wks and 4wks Klf9KI hearts compared to Wt-Cre. These data indicate progressive mitochondrial dysfunction with constitutive Klf9 expression, which results in progressive cardiac hypertrophy, dysfunction leading to the development of heart failure and death.

**Figure 4.**
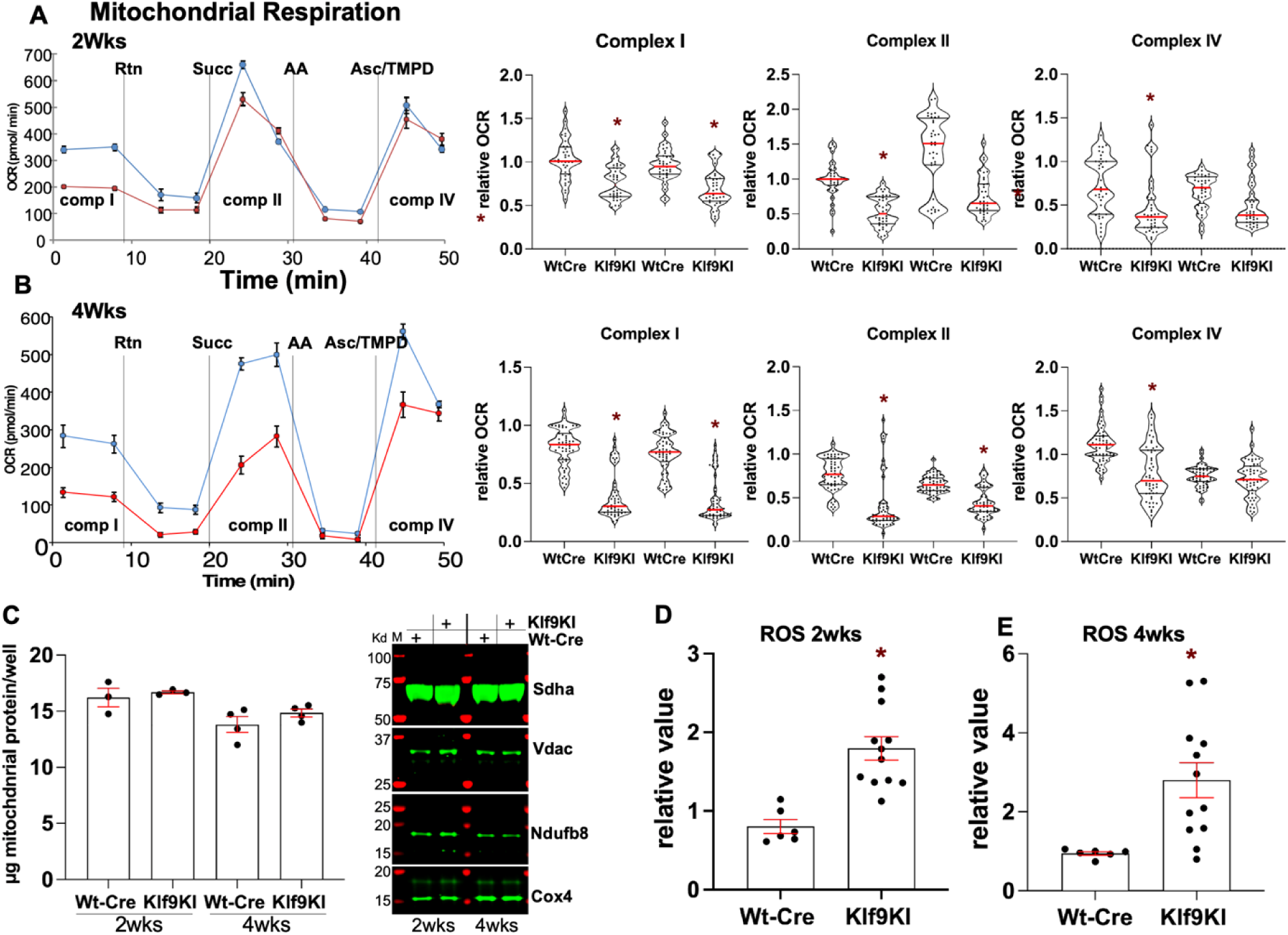
Mitochondrial function and oxidative stress in Klf9KI versus Wt-Cre hearts. **A** and **B.** Mitochondria were freshly isolated from the hearts of 2wks **(A)** and 4wks **(B)** old mice and plated in mitochondrial assay buffer containing pyruvate, malate, and FCCP. The OCR (pmol/min, y axis) over time (min, x axis) was measured using the Seahorse XFe96 analyzer, before and after the sequential injections of rotenone (Rtn), succinate (succ), antimycin A (AA), and Ascorbic acid + TMPD (Asc/TMPD), at 2 time points for each injection, as seen on curve. Violin plots on the right show the relative OCRs at both the time points for each injection graphed separately for Basal (Complex I), Complex II, and Complex IV mediated respiration. The data were normalized to mitochondrial protein per well. **C.** Left; graph showing µg of mitochondria/well for Seahorse experiments in (A) and (B). Right; representative western blots with mitochondrial proteins showing equal loading of samples. **D.** Graphs show hydrogen peroxide production as a measure of ROS to determine oxidative stress in hearts of 2wks and 4wks old Wt-Cre and Klf9-KI mice. For all graphs, error bars represent SEM, * is p<0.05 compared to respective Wt-Cre, N=3-5 independent hearts.

## Discussion

Heart endures constant workload that requires steady supply of energy, additionally, it has the capacity to significantly increase the work output depending on the demands in response to both physiological and pathological conditions (7). Thus, for cardiomyocyte integrity it is essential to maintain contractile and metabolic homeostasis, the two highly interdependent factors required for heart function. Heart being an omnivore, can utilize a variety of substrates depending on availability and demands, which requires adaptation of gene expression and signaling pathways, metabolite concentrations and substrate availability. High glycolytic flux in the neonate hearts switches to mainly fatty acid oxidation for energy in adult hearts. (8). We recently reported that Klf9 regulates GR dependent metabolic adaptations in cardiomyocytes. We identified and reported the genomic targets of Klf9 in Dex treated neonatal cardiomyocytes, and by examining the transcriptome with knockdown of Klf9, validated GR-Klf9 mediated regulation of these metabolic genes and its effects on glycolysis and mitochondrial function (3). We reported Klf9 as one of the early direct targets of GR in cardiomyocytes (9). Here we show that cardiac Klf9 expression decreases during compensatory cardiac hypertrophy, when mice are subjected to pressure overload, suggesting that a decrease in Klf9 levels could be contributing to pathogenesis. These data align with recent publication by Zhang et. al. that shows similar decrease in Klf9 expression in hearts undergoing Ang II induced cardiac hypertrophy and failure. The authors report that Klf9 through its actions on PGC-1α and Mfn2 regulates mitophagy and mitochondrial function. Inducible, conditional Klf9 transgenic mice rescued Ang II induced cardiac dysfunction, hypertrophy and fibrosis. On the other hand, conditional Klf9 knockout exaggerated the Ang II induced pathological hypertrophy, with early onset of heart failure (4). Meanwhile, we were characterizing Klf9KI mice, mated with αMHC-Cre, for role of Klf9 in postnatal heart development, and to examine its sufficiency in cardiac metabolic adaptations under basal and induced cardiac stress conditions. However, our data shows that constitutive Klf9 expression result in the development of spontaneous metabolic and cardiac dysfunction by 4wks of age, failure by 8wks and death by 12-14wks of age. These differences could be due to the inducible versus constitutive nature of the Klf9 expression, where our Cre and hence Klf9 expression could be as early as E10.5-11.5 (10) (11), and exogenous Klf9 could be interfering with the metabolic switch from glycolysis to fatty acids with postnatal development. However, independent cardiac studies have also reported a detrimental role of Klf9 in cardiomyocytes with ischemic injury (12), during streptozotocin-induced diabetic cardiomyopathy (13) and inflammatory injury with myocardial ischemia (14).

Klf family members share a conserved C-terminal DNA binding domain, however, the variable regulatory N-terminal domain based on its associating factors and can have different effects on target gene expression. Based on structural domains and association with corepressor Sin3A, Klf9 has been shown function as transcriptional repressor (1) (15) (16). Interestingly, our data from 1wk old Klf9KI hearts aligns with the predicted transcriptional repressive function of Klf9, where we report a downregulation of ∼64% of the differentially expressed genes compared to Wt-Cre. At 4wks and 8wks, we see significant increase in genes involved in innate immunity, apoptosis and extracellular matrix, which we believe are secondary to the ensuing pathogenesis and independent of Klf9 targeting, in addition to the changes we observe in cellular signaling, organelle structure and morphology with later time points. Notably, significant increase in 3-hydroxybutyrate dehydrogenase 1 (Bdh1) expression is observed starting P1, suggesting utilization of ketone metabolism at an early age, which could be due to compromised glucose and/or lipid metabolism in the Klf9KI neonate hearts, compared to Wt-Cre. Our transcriptome data from 1wk Klf9KI hearts show decrease in genes of carbohydrate and glycogen metabolism pathway (Fbp2, Pygl), a stage where glucose serves as the main source of energy compared to other substrates. Bdh1, expressed in adult hearts with little to no expression in neonates (17) increases during heart failure in both humans and rodents (18) (19) (20). Cardiac ketone flux serves as an alternative, compensatory measure by the heart during metabolic crisis, where it supports metabolic remodeling with intermittent fasting or exercise in striated muscle (21) or decrease oxidative stress and cardiac dysfunction with pressure overload induced heart failure (22).

Our data with the untargeted metabolome revealed novel insights on the differential accumulation of the metabolites in the Klf9KI compared to Wt-Cre hearts. We chose untargeted approach to identify as many differential tissue metabolites as possible and correlate with the observed hypertrophic phenotype seen in the Klf9KI mice (23). Integrating the metabolome data with RNAseq data from 4wks hearts of Klf9KI and Wt-Cre mice, showed strong correlation of the differential transcriptome and its effects on the metabolome. Now, based on these results we can have a more targeted approach and in future characterization of these hearts we intend to include targeted metabolite analysis including lipid, carbohydrates and amino acids metabolism. An interesting finding was that although, complex I activity was decreased at both 2 and 4wks of age, a compromised complex II and complex IV was seen only at 4wks in Klf9KI mitochondria compared to Wt-cre. It has been shown that mitochondrial complex II is the source for respiratory reserve capacity (24), which plays an essential role in accommodating supply to increasing demands (25) (26). We hypothesize that the even with decreased complex I, compensated respiratory reserve capacity could be maintaining energy supply in the Klf9KI hearts to avert ATP crisis, due to metabolic reprogramming with constitutive expression of postnatal Klf9.

Thus, the study highlights a central role of Klf9 in maintaining the cardiac metabolic homeostasis. Klf9 expression and function could be under control of external stimulus like circulating corticosteroids (9) or upstream still unidentified signaling pathways that dictates the expression level and chromatin association of endogenous Klf9 for fine tuning of the metabolic adaptations to changing physiological variations and pathological conditions.

## Author contributions

CT performed experiments; SA assisted in the animal colony, genotyping and initial generation of the Klf9KI mice; AV assisted in microscopy of histology sections; ZY performed the sham/TAC surgeries and Echo on the mice; NS assisted in RNA seq data analysis; MA assisted in the Seahorse experiments, DS designed experiments, performed data analysis with figures and wrote the manuscript.

## Acknowledgements

We thank Dr. Sadoshima, Chair of department of Cell Biology and Molecular Medicine, Rutgers New Jersey Medical School for support.

## Materials and methods

### Animals

All animal procedures in this study were performed in accordance with the NIH Guidelines for the Care and Use of Laboratory Animals (National Academies Press, No. 85-23, 2011). The animals were housed in Animal facility at Rutgers, The State University of New Jersey located in Newark NJ, as per standard procedures/protocols. All protocols were approved by the IACUC of the Rutgers-New Jersey Medical School. C57BL/6 mice for Klf9-ChIP-Seq and Sham and transverse aortic constriction (TAC) surgeries were purchased from The Jackson Laboratory. Cardiac-specific Klf9 transgenic mice were generated by crossing ROSA26_loxP_αMHC_Klf9_KnockIn mice (generated at Cyagen) with αMHC-Cre mice.

### Transverse Aortic Constriction

TAC was performed as described previously [1, 2]. Briefly, a 7-0 braided polyester suture was tied around the transverse thoracic aorta, against a 27 gauge needle, between the innominate artery and the left common carotid artery. Control mice were subjected to a sham operation involving the same procedure, minus the aortic constriction.

### Survival analysis

Cohorts of 26 Klf9 KI mice (12 Male and 14 Female) along with age and sex-matched Wt Cre were recruited and observed daily to record their survival. Event occurred was entered as “1” when mice were found dead while a “0” was entered when the mice were found healthy at the endpoint of observation. Survival analyses were performed using the Kaplan-Meier survival analysis plot in GraphPad Prism 10.0.2.

### Echocardiography

Echocardiographic measurements were performed as described previously [1, 2]. Briefly, mice were anaesthetized with Ketamine (80-100 mg/kg)/Xylazine (10mg/kg) administered by intraperitoneal injection. The adequacy of the anesthetic was confirmed by the tail pinch. Transthoracic echocardiography was performed using the Vevo 3100 imaging system (Visual Sonics) with a MX400 30 MHz (mouse, cardiology) scan head encapsulated transducer. Electrocardiographic electrodes were taped to the 4 paws, and then 1D M-mode and 2D B-mode tracings were recorded from the parasternal short-axis view at the mid-papillary muscle level. Vevo 3100 Software (Vevo Lab, version 3.2.6) was used for image capture and analysis.

### ChIP**-**Seq

Chromatin immunoprecipitation sequencing (ChIP-Seq) was performed as reported earlier [3]. Hearts from 12week-old mice were harvested, snap frozen in liquid nitrogen and sent to Active Motif (Carlsbad, CA) for Klf9-ChIP-Seq. Immunoprecipitation was performed using anti-KLF9 antibody (A7196, ABclonal, Inc., Woburn, MA) followed by high throughput Illumina NextSeq 500 sequencing. Input DNA was taken before immunoprecipitation and served as a control for normalization and eliminating background.

### ChIP**-**Seq Data Analysis

Bioinformatics on the sequencing data generated was performed by Active Motif, Inc., and done as described previously [3]. The Data explanation from Active Motif includes the following: Sequence analysis – 75-nt sequence generated by sequencing reads were mapped to the genome using BWA algorithm, and the information stored in BAM format. Only reads which align with no more than two mismatches and map uniquely are used in analysis. Determination of Fragment Density – Aligned reads (tags) are extended in silico at their 3′ ends to a length of 200 bp using Active Motif software, corresponding to the average fragment length in the size-selected library. To identify the density of fragments along the genome, the genome is divided into 32-nt bins and the number of fragments in each bin is determined and stored in BigWig file and can be visualized on genome browsers. Peak Finding – Genomic regions with enrichments in tag numbers are termed ‘intervals’, and defined by the chromosome number, start and end coordinate. 24237848 normalized tags were used for peak calling using MACS 2.1.0 algorithm with default cutoff p value 1e-7. Peak filtering was performed by removing false ChIP-Seq peaks as defined within the ENCODE blacklist. Merged region analysis – To compare peak metrics between two samples, overlapping Intervals are grouped into ‘Merged Regions’, which are defined by start coordinate of the most upstream interval and end coordinate of the most downstream interval (=union of overlapping intervals; “merged peaks”). In locations where only one sample has an interval, this interval defines Merged Region. The use of Merged Regions is necessary because the locations and lengths of intervals are rarely exactly same when comparing different samples. Annotations – Intervals, Merged Regions, their genomic locations along with proximities to gene annotations and other genomic features are determined and presented in Excel spreadsheets. Average and peak fragment densities within Intervals and Active Regions are compiled.

The Merged Regions provides the peak metrics for samples in all peak regions and used to analyze ChIP enrichments in the samples. The MaxTags filter out the regions with low signal values, which are expected to be more variable and therefore lead to unreliable, often exaggerated ratios. To calculate the maximum number of tags (reads) multiply the average value of the tag per sample with the length of the regions and divide by constant 224. The 224 value represents the average length of the sequenced fragment in base pairs. As the signal is normalized in 32 bp bins along the genome, 7 bins of 32 bp length (224) is equivalent to the fragment length of 200 bp. Cutoffs were set at 50 MaxTags for Klf9-ChIP-seq. From the sorted data, subsets were made for manuscript preparation with proper data representation and presentation.

### RNA Sequencing

Hearts from 1wk, 4wks, and 8 wks old Klf9KI male mice and their corresponding Wt Cre controls were collected and sent to Novogene Corporation Inc for RNAseq. Briefly, mRNA was purified from total RNA using poly-T oligo-attached magnetic beads. After fragmentation, the first strand cDNA was synthesized using random hexamer primers, followed by the second strand cDNA synthesis using either dTTP for non-strand specific library or dUTP for strand specific library. These libraries were ready after end repair, A-tailing, adapter ligation, size selection, amplification, and purification. The libraries were checked with Qubit and real-time PCR for quantification and bioanalyzer for size distribution detection. After library quality control, different libraries were pooled based on the effective concentration and targeted data amount, then subjected to Illumina sequencing. Sequenced reads were processed through fastp software to remove possible adapter sequences and nucleotides with poor quality to obtain clean reads. Paired-end clean reads were aligned to the reference genome using Hisat2 v2.0.5. Alignment information for each read was stored in the BAM format. The mapped reads of each sample were assembled by StringTie (v1.3.3b) in a reference-based approach. featureCounts v1.5.0-p3 was used to count the reads numbers mapped to each gene. Distribution of read counts in libraries was examined before and after normalization. The original read counts were normalized to adjust for various factors such as variations of sequencing yield between samples. These normalized read counts were used to accurately determine differentially expressed genes using DESeq2 R package (1.20.0). Log2 fold change between the groups was calculated and the Wald test was used to determine statistical significance (p-value). Adjusted p values were calculated using Benjamini-Hochberg procedure. Genes with an adjusted padj<0.05 and log2FC of at least 1 were called out as differentially expressed genes for the comparison between the groups. Heatmap for the differentially regulated genes was generated using Heatmapper [4], with average linkage for clustering.

### Genome browsers

Integrated Genome Browser (IGB) [5] and Integrated Genomic Viewer (IGV) [6] was used for visualization of Klf9-ChIP-seq and RNAseq data, respectively.

### Functional annotation and GO terms

Functional annotation was performed using by Database for Annotation, Visualization and Integrated Discovery (DAVID) algorithm [7].

### Metabolomic Analysis

Hearts from 4wks old Wt Cre and Klf9 KI mice were collected and sent to Creative Proteomics for untargeted metabolomics. The sample preparation and analyses from Creative Proteomics includes the following–Samples were thawed on ice and methanol at 8μL/mg the raw material and two 5-mm metal balls were added to the tube. All samples were ground 180 s at 65 Hz, twice, followed by sonication for 30 min, 4°C. Then each sample was kept at -20°C for 1 h, vortexed for 30 s, and centrifuged 10 min at 12,000 rpm, 4°C. Finally, 200μL of supernatant was transferred and 5μL of 0.14 mg/mL DL-o-Chlorophenylalanine was added into vial followed by filtration through a 0.22μm filter for LC-MS analysis. Separation was performed by Vanquish Flex UPLC combined with Q Exactive plus (Thermo) and screened with ESI-MS to analyze the metabolic profiling in both ESI positive and ESI negative ion modes. The raw data are acquired and aligned using the Compound Discoverer (3.0, Thermo) based on the m/z value and the retention time of the ion signals. Ions from both ESI- or ESI+ are merged and imported into the SIMCA-P program (version 14.1) for multivariate analysis. A PCA is first used as an unsupervised method for data visualization and outlier identification. Supervised regression modeling is then performed on the data set by use of Partial Least Squares Discriminant Analysis (PLS-DA) or Orthogonal Partial Least Squares Discriminant Analysis (OPLS-DA) to identify the potential biomarkers. The biomarkers are filtered and confirmed by combining the results of the VIP values (VIP > 1.5), t-test (p < 0.05) and fold change values (|log2FC| > 0.5). The chemical structures of important metabolites were then identified according to online databases such as the Human

Metabolome Database (www.hmdb.ca) using the data of accurate masses and MS/MS fragments. When necessary, further confirmation was acquired through comparisons with authentic standards, including retention times and MS/MS fragmentation patterns. Hierarchical cluster analysis of metabolome data from significant metabolites was performed using the complete linkage algorithm of the program Cluster 3.0 (Stanford University) and the results are visualized using Pheatmap 1.0.12 (Raivo Kolde). Pathway analysis of significant metabolites was performed using MetaboAnalyst.

### Western Blotting

Total protein lysates were isolated in RIPA buffer (Thermo Scientific) containing Halt protease inhibitor (PI) cocktail (Thermo Scientific). Mitochondrial protein was isolated as described in the mitochondria isolation section. Protein was quantified by BCA protein assay kit (Thermo Scientific), separated on gradient (4%–12% gel) XT gels (Biorad), and transferred to 0.2 μm nitrocellulose membrane. Membranes were probed with primary antibodies for KLF9 (Abclonal#A7196), Myh7 (Sigma#M8421), Lamin A/C (Santacruz#SC-376248), BDH1 (Proteintech#15417-1-AP), HMGCS2 (Cell Signaling Technology#20940s), VDAC (Cell Signaling Technology#4866s [tissue lysates]; #4661 [mitochondrial protein]), SDHA (Cell Signaling Technology#11998), Ndufb8 (Abcam#EPR15961), COX4 (Cell Signaling Technology#4850), GAPDH (Cell Signaling Technology#97166s), ANKRD1 (CARP) (Santacruz#sc-365056), and Nppa (ANP) (Invitrogen#702539) in 5% BSA blocking buffer. The Western blot signals were detected by the Odyssey imaging system (LI-COR).

### Quantitative PCR

Total RNA was isolated from cardiomyocytes using TRIzol reagent (Life Technologies) and was reverse-transcribed to cDNA using a High-Capacity cDNA Reverse Transcription Kit (Thermo Fisher Scientific), as per manufacturer’s protocol. The cDNA was used for qPCR with an Applied Biosystems 7500 thermocycler using Taqman gene expression assays. The following Taqman primers were used for qPCR analysis: Klf9 (ID: Mm00495172_m1), Acta1 (ID: Mm00808218_g1), Myh7 (ID: Mm00600555_m1), Klf10 (ID: Mm0449812_m1). 18S (ID: mm03928990_g1) was used as internal control for normalization.

### Mitochondria isolation and electron flow assay

Mitochondria were isolated from cardiac tissue using differential centrifugation as described previously [8] with modifications as required. Briefly, freshly isolated hearts were rinsed in ice-cold PBS followed by rinsing, chopping, and homogenization in ice-cold mitochondrial isolation buffer (70 mM sucrose, 210 mM mannitol, 5.0 mM HEPES, 1.0 mM EGTA, and 0.5 % (w/v) fatty acid–free BSA, pH 7.0) containing PI cocktail. The homogenate was centrifuged at 800g for 10 mins at 4°C to remove debris and extremely large cellular organelles followed by centrifugation at 8000g at 4°C for 10 mins. The isolated mitochondrial pellet was reconstituted in ice-cold mitochondrial isolation buffer (containing PI cocktail) followed by dilution and plating in mitochondrial assay buffer (70 mM sucrose, 220 mM mannitol, 10 mM KH2PO4, 5 mM MgCl2, 2 mM HEPES, 1.0 mM EGTA, 0.2 % (w/v) fatty acid–free BSA, pH 7.2) containing 10 mM pyruvate, 2 mM malate, and 4 μM FCCP. Using the Seahorse XFe96 Analyzer, the oxygen consumption rate (OCR) was measured at baseline and after sequential injections of 2 μM rotenone, 10 mM succinate, 4 μM antimycin A, and 10 mM ascorbate plus 100 μM tetramethyl-p-phenylene diamine (TMPD) at 37°C. The OCR values were normalized to mitochondrial protein concentration.

### Oxidative stress

Hydrogen peroxide (H_2_O_2_) as a measure of oxidative stress was detected using Amplex Red H_2_O_2_ assay kit (Invitrogen) as described previously [9] with modifications as required. Briefly, small sections of harvested heart tissue were incubated with Amplex Red (100 uM) and horseradish peroxidase (1 U/mL) for 30 min at 37°C in Krebs-Hepes buffer protected from light. The supernatant was collected, and fluorescence was measured alongside H_2_O_2_ standards using 540 nm excitation and 590 nm emission filters. H_2_O_2_ release from the tissue sections was calculated using the standard curve and normalized to tissue weight.

### Histological analysis

Hearts were fixed with 10% formalin and embedded in paraffin. Paraffin-embedded tissue slices (5 um) on slides were deparaffinized, rehydrated and stained with Picric Acid Sirius Red (fibrosis), Wheat Germ Agglutinin (cell size), and Hematoxylin & Eosin (tissue structure and cell morphology) stains. Tissue-slices were washed, dehydrated, and mounted in permanent mounting medium.

### Statistics

Klf9-ChIP-Seq statistics (provided by Active Motif) and RNA-Seq statistics (provided by Novogene) and metabolomics statistics (provided by Creative Proteomics) is shown in detail in the respective sections. Calculation of significance between 2 groups was performed using an unpaired, 2-tailed Students t-test). All experiments conducted as minimum N=3 and presented as average with SEM, p<0.05 was considered significant.

