## Supplementary Figures for "Constitutive expression of cardiomyocyte Klf9 precipitates metabolic dysfunction and spontaneous heart failure"

[†] To whom correspondence should be addressed:

**Figure S1.**

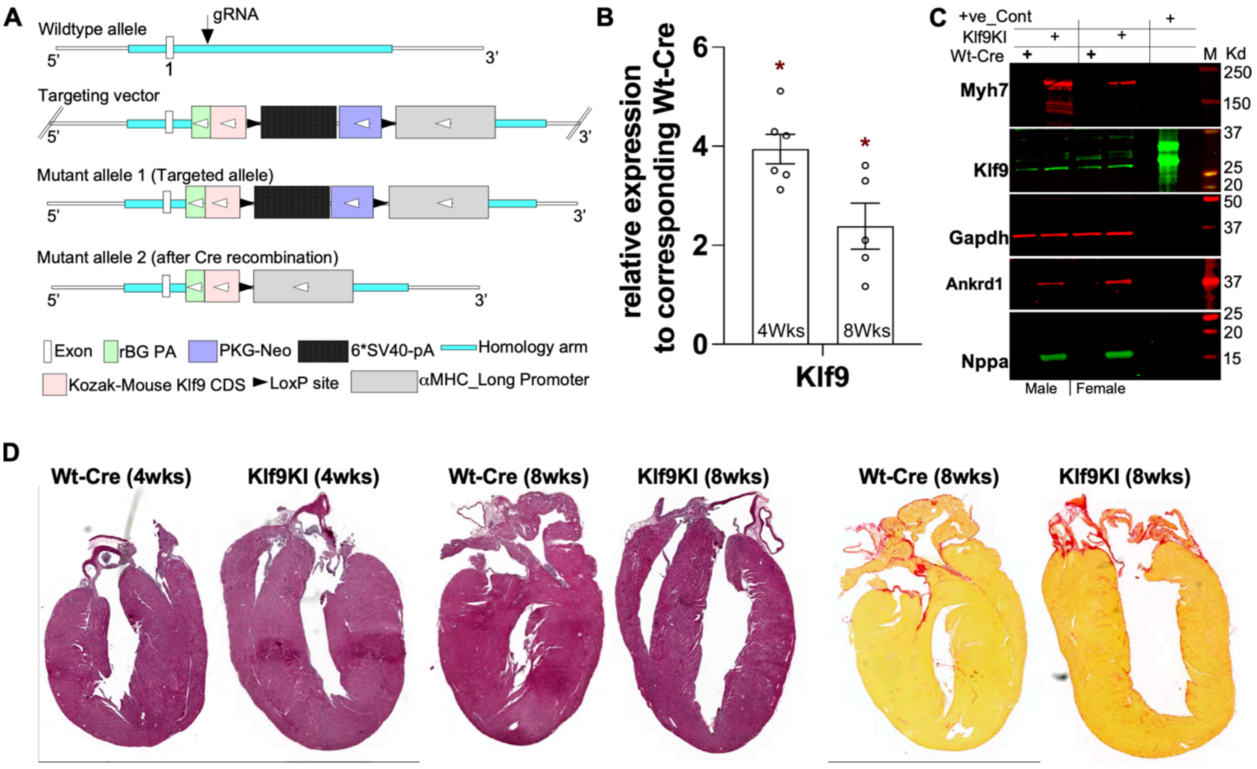


**Figure. S1.** Cardiac-specific Klf9KI mice. **A.** Mouse Klf9 CDS with αMHC promoter was inserted into ROSA26 locus with flaking loxP sites for conditional expression with Cre. **B**. Graph represents relative Klf9 transcript abundance in Klf9KI compared to Wt-Cre hearts in 4wks and 8wks mice. Error bars represent SEM, * is $p<0.05$, n= 5-6. **C.** Western blot of total heat lysate from male and female Klf9 and respective Wt-Cre mice for indicated proteins Ankrd1 and Myh7 shown as markers for cardiac hypertrophy. Gapdh shown for loading control. **D**. Histological analyses on longitudinal sections from hearts of adult mice (4-8 weeks), left, H & E staining, right PASR staining for fibrosis.

### Figure S2

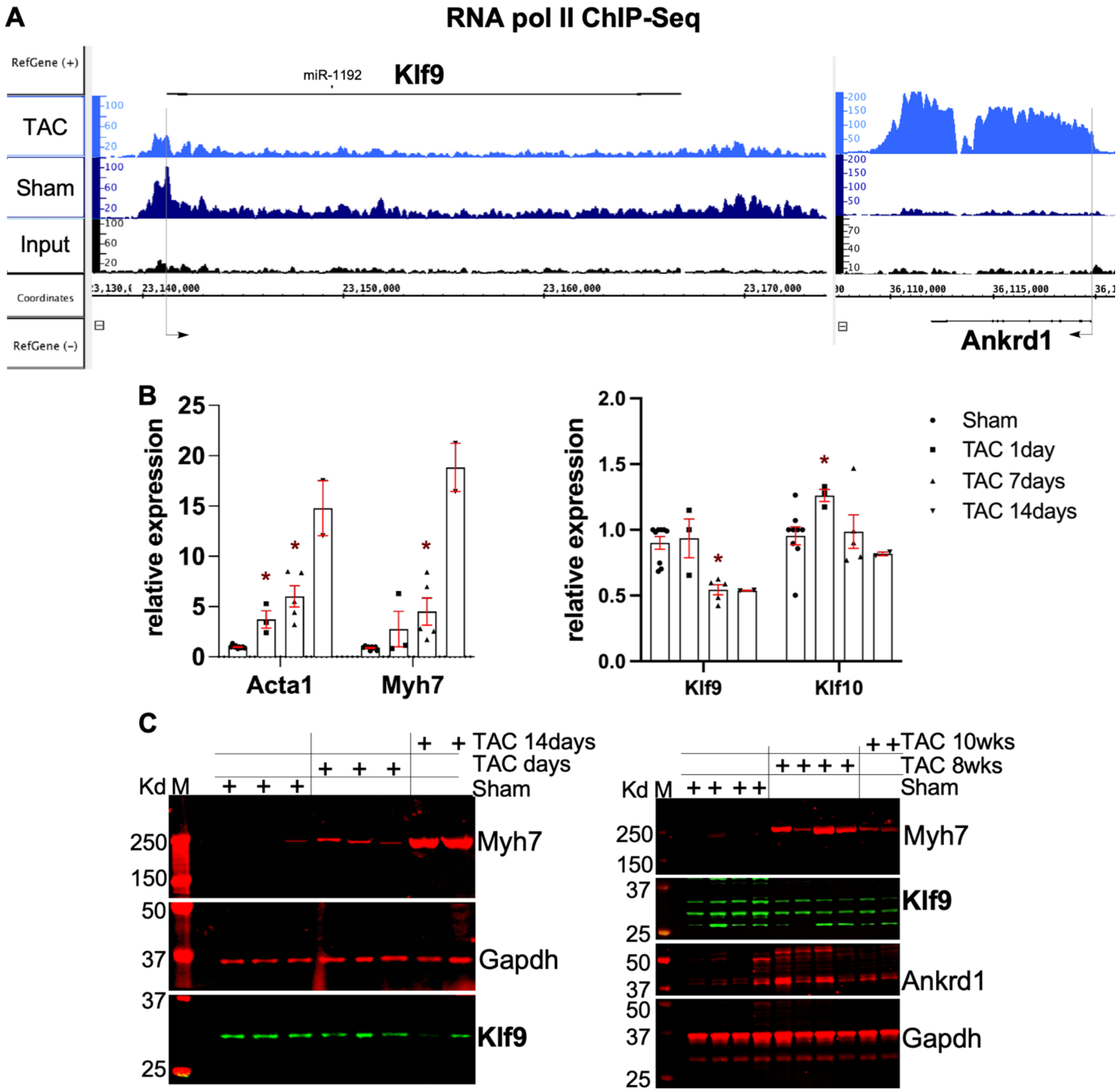


**Figure S2.** Cardiac Klf9 expression in pressure overload induced cardiac hypertrophy. **A.** Plots shows RNA pol II occupancy and distribution across the Klf9 and Ankrd1 gene structure in Sham and TAC (4-7days) hearts, along with Input. X' represents chromosomal coordinates, gene structure, arrow shows direction of transcription from promoter regions. Y' shows the signal tracks and fragment densities of RNA pol II (GSE50637). **B.** Graph represents qPCR data from Sham/TAC hearts for indicated time periods of 1day, 7days, 14days. Klf10 is shown as comparative gene for Klf9. Acta1 and Myh7 are shown as markers for progressive hypertrophy. Error bars represent SEM, * is $p<0.05$ compared to respective Sham, N=3-5. **C.** Western blots on protein lysates from Sham/TAC hearts with indicated time periods of 7days, 14days, 8wks or 10wks.

**Figure S3**

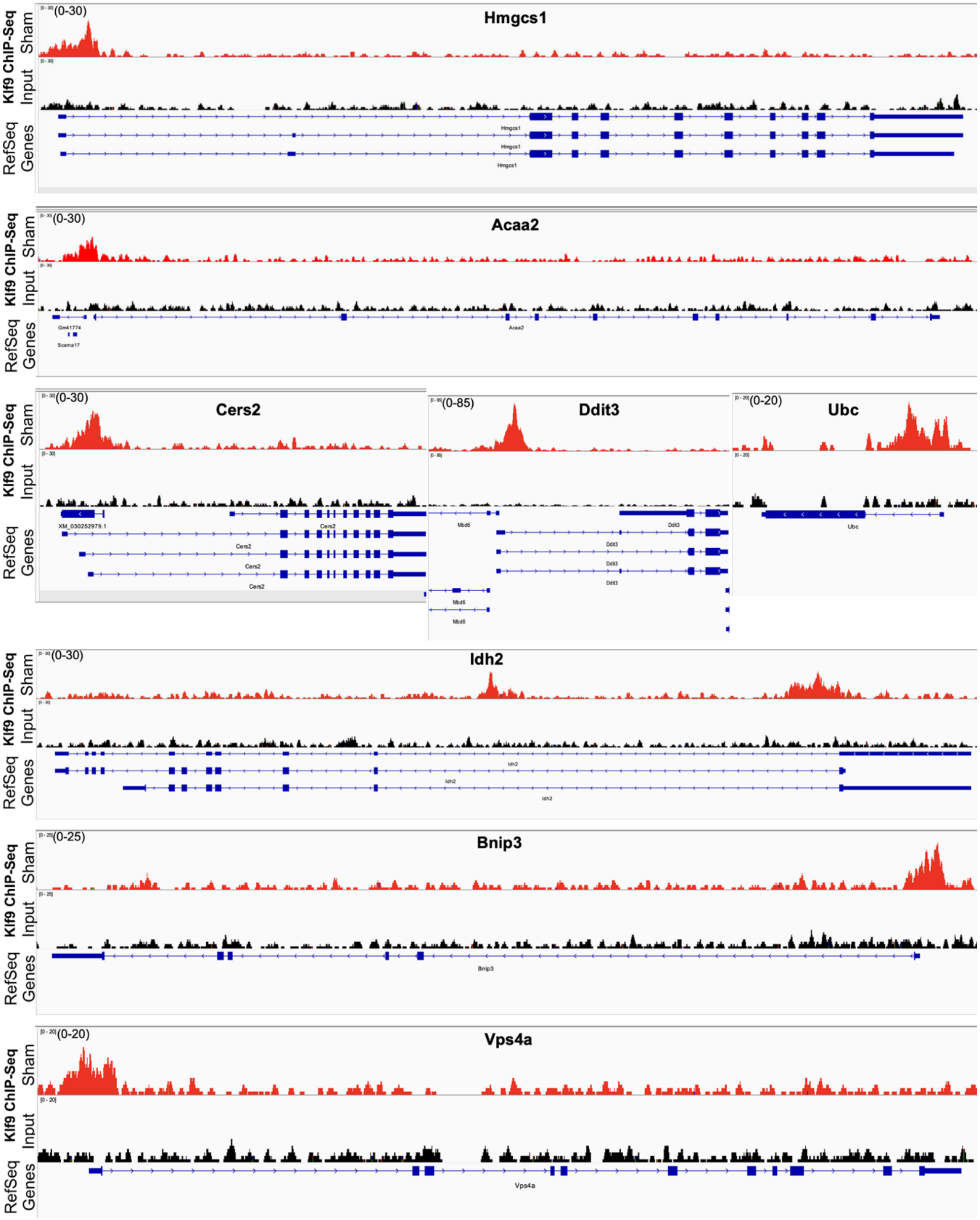


**Figure S3.** Klf9 occupancy on representative genes in adult mouse hearts. Plots show KLf9 occupancy on adult (sham) hearts and input samples, as viewed on integrated genome browser. The X axis represents chromosomal coordinates, reference sequence and gene structure, Y axis shows the peak value on the signal tracks of Klf9 genomic occupancy-.

**Figure S4**

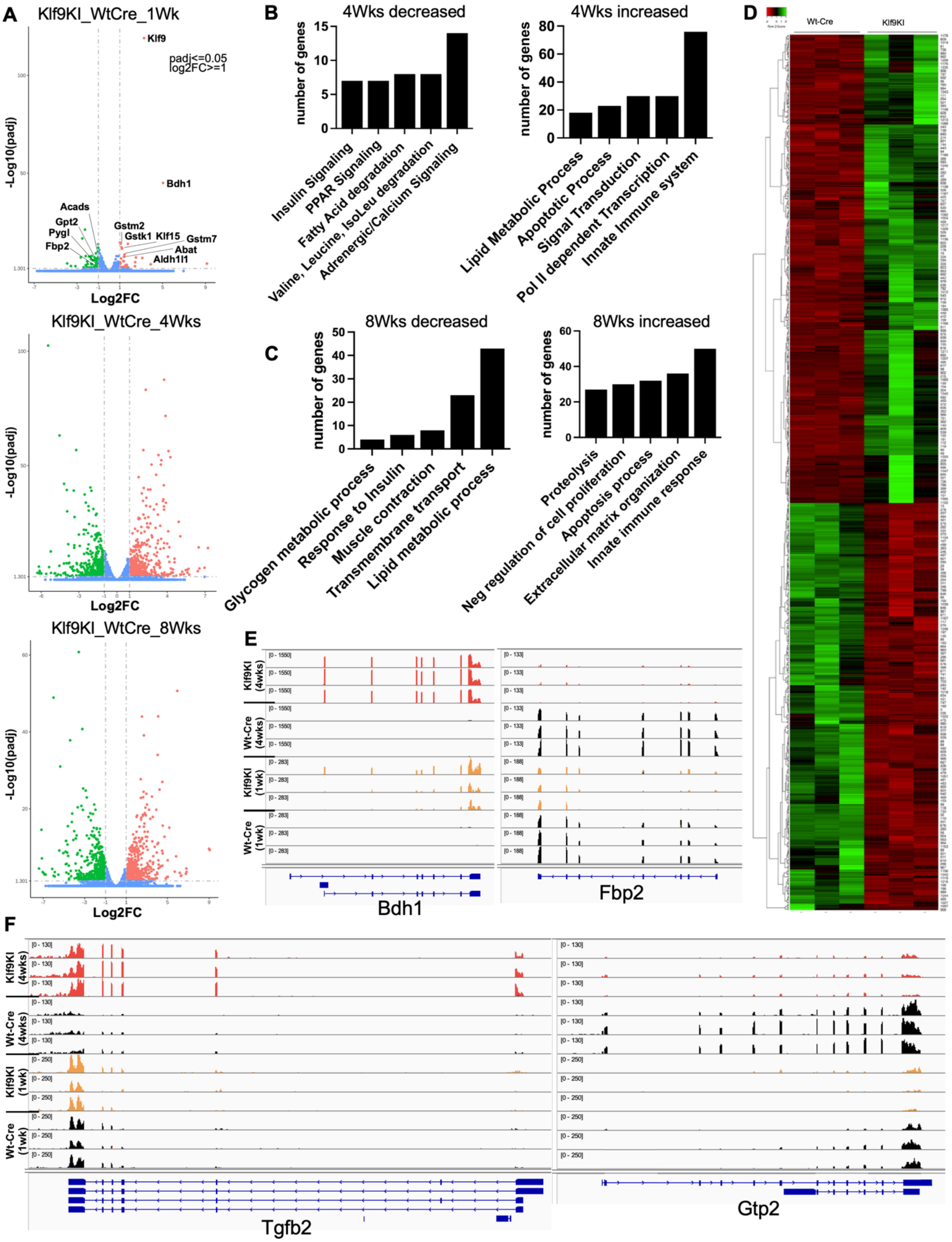
A
Klf9KI_WtCre_1Wk
padj<=0.05
log2FC>=1
Klf9
Bdh1
Acads
Gpt2
Pygl
Fbp2
Gstm2
Gstk1
Klf15
Gstm7
Abat
Aldh1l1
-Log10(padj)
Log2FC
Klf9KI_WtCre_4Wks
Klf9KI_WtCre_8Wks
B
4Wks decreased
number of genes
Insulin Signaling
PPAR Signaling
Fatty Acid degradation
Valine, Leucine, IsoLeu degradation
Adrenergic/Calcium Signaling
4Wks increased
Lipid Metabolic Process
Apoptotic Process
Signal Transduction
Pol II dependent Transcription
Innate Immune system
C
8Wks decreased
Glycogen metabolic process
Response to Insulin
Muscle contraction
Transmembrane transport
Lipid metabolic process
8Wks increased
Proteolysis
Neg regulation of cell proliferation
Apoptosis process
Extracellular matrix organization
Innate immune response
D
Wt-Cre
Klf9KI
E
Klf9KI (4wks)
Wt-Cre (4wks)
Klf9KI (1wk)
Wt-Cre (1wk)
Bdh1
Fbp2
F
Tgfb2
Gtp2

**Fig S4. A.** Volcano plots showing genes that are upregulated (red), downregulated (green), and unchanged (blue) between Wt-Cre and Klf9KI mice in 1wk, 4wks and 8wks Klf9KI hearts compared to Wt-Cre. **B.** Graphs lists the number of genes in the functional groups as categorized by KEGG pathway analysis, which show differential regulation in 4wks hearts of Klf9KI compared to Wt-Cre. **C**. Graphs lists the number of genes in the functional groups as categorized by KEGG pathway analysis, which show differential regulation in 8wks hearts of Klf9KI compared to Wt-Cre. **D.** Heatmap displays the 1247 genes that are differentially regulated in Wt-Cre and Klf9KI mice at 8wks. **E** and **F.** Screenshot of the Integrated genome Browser showing alignment of the RNAseq data of representative genes from Wt-Cre and Klf9KI mice at 1wk and 4wks on the reference genome. X axis represents ref sequence and gene structure, while Y axis shows the values on the signal tracks. The value is kept constant across the samples for each gene. RNAseq data included n=3 for each group.
